# Traumatic brain injury exacerbates mitochondrial dysfunction in APP/PS1 knock-in mice through time-dependent pathways

**DOI:** 10.1101/2025.10.01.679790

**Authors:** Elika Z. Moallem, Hemendra J. Vekaria, Teresa Macheda, Margaret R. Hawkins, Kelly N. Roberts, Samir P. Patel, Patrick G. Sullivan, Adam D. Bachstetter

## Abstract

Cerebral hypometabolism occurs in both traumatic brain injury (TBI) and Alzheimer’s disease (AD), but whether these conditions act through distinct or overlapping mechanisms is unclear. TBI disrupts cerebral metabolism via blood–brain barrier damage, altered glucose transporter expression, calcium buffering abnormalities, and oxidative damage to metabolic enzymes. AD-related hypometabolism is linked to amyloid-β (Aβ) effects on mitochondria, including impaired respiration, oxidative stress, and altered mitophagy, fusion, and fission. We tested whether TBI-induced mitochondrial dysfunction exacerbates Aβ-mediated impairment using a closed-head injury (CHI) model in APP/PS1 knock-in (KI) mice. Injuries were delivered at 4-5 months of age, before plaque formation and mitochondrial deficits in KI mice. Bioenergetics were measured at 1, 4, and 8 months post-injury in hippocampus and cortex using Seahorse assays on isolated mitochondria. At 1 month, genotype-by-injury interactions revealed greater dysfunction in KI mice than either condition alone, with males more vulnerable than females. At 4-8 months, amyloid-mediated effects predominated, while TBI-specific changes were no longer apparent, suggesting recovery or convergence onto shared mechanisms. These results indicate that TBI can temporarily worsen mitochondrial dysfunction in the context of early amyloidosis, with sex influencing vulnerability. Findings provide insight into the temporal relationship between TBI and amyloid-induced mitochondrial deficits and support the importance of sex as a biological variable in neurodegenerative disease progression.

**Highlights:** TBI and APP/PS1 genotype interact to worsen early mitochondrial dysfunction.

Hippocampus exhibits greater susceptibility to combined TBI and amyloid pathology.

Sex-specific effects and temporal patterns underscore TBI’s role in AD risk.

Male mice show heightened vulnerability to TBI-induced mitochondrial dysfunction.

**GRAPHICAL ABSTRACT:** 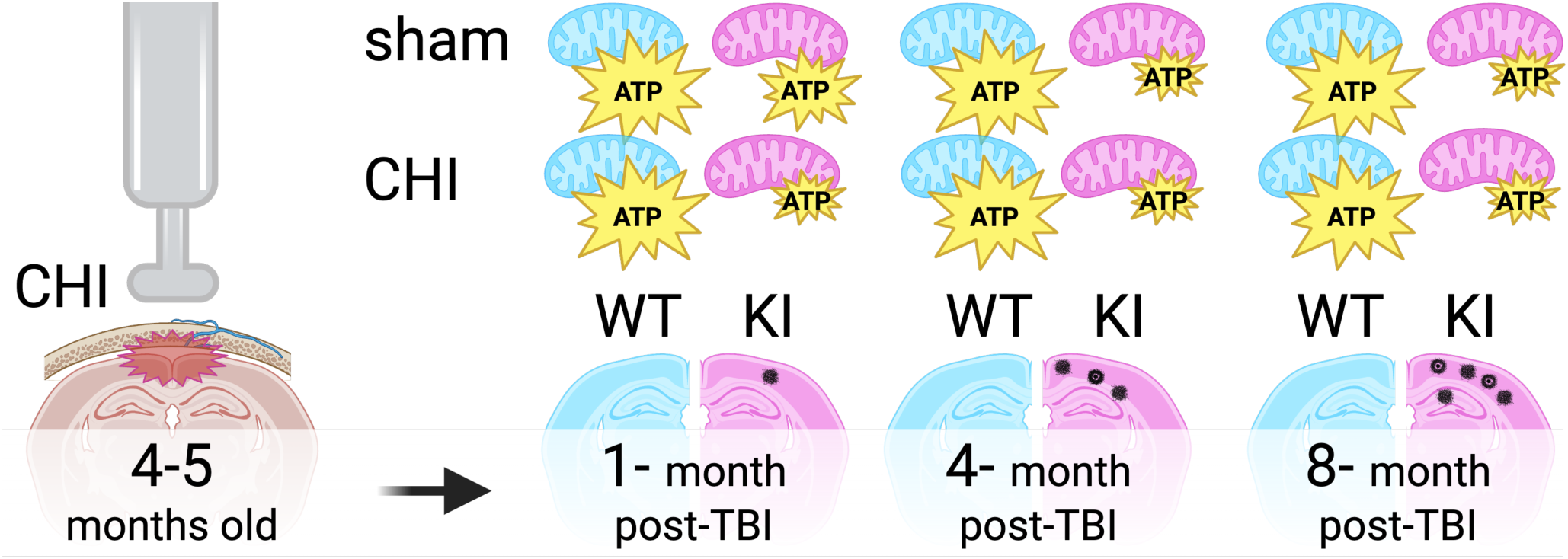

## Introduction

Traumatic brain injury (TBI) is an environmental risk factor for Alzheimer’s disease (AD), with human population-level data showing a strong association between TBI and subsequent dementia development [1–4]. Mitochondrial dysfunction plays a central role in both conditions, contributing to cerebral hypometabolism, neuronal energy deficits, and oxidative stress in people and experimental models [5–7]. In experimental TBI models, an initial compensatory hyperglycolysis and heightened oxygen utilization near the lesion site is typically followed by prolonged hypometabolism, with metabolic shifts including increased flux through the pentose phosphate pathway (PPP) to support DNA repair processes [8, 9]. These compensatory mechanisms are often insufficient to prevent long-term bioenergetic deficits, as evidenced by hyperglycemia and increased plasma glucose levels in severe TBI patients [10, 11]. Studies using Seahorse and spectrophotometric assay measurements in experimental models have demonstrated that TBI and aging disrupts mitochondrial respiration, reducing adenosine triphosphate (ATP) production and impairing electron transport chain (ETC) function, particularly Complexes I, II and IV [12–15]. Mitochondria isolated from injured brains in controlled cortical impact (CCI) models display significant declines in oxidative phosphorylation capacity following CCI [16–18]. Similarly, AD exhibits reduced brain glucose metabolism, particularly in temporal and parietal regions in people [19, 20], with animal models demonstrating decreased glucose utilization in the hippocampus and cortex [21–24]. Several studies in experimental models have demonstrated that amyloid precursor protein (APP) and amyloid beta (Aβ) can localize to mitochondria, influencing mitochondrial dynamics and respiratory function [25–28]. Furthermore, experimental model studies show that γ-secretase complexes within mitochondria have been shown to generate Aβ locally, suggesting mitochondrial dysfunction may directly contribute to intramitochondrial Aβ production [28, 29]. Mouse models of cerebral amyloidosis exhibit early mitochondrial abnormalities, including increased mitophagy, reduced mitochondrial biogenesis, and decreased ATP production [30–32]. Given these parallel metabolic deficits, TBI may potentially accelerate mitochondrial failure in those susceptible to Aβ pathology.

The aim of this study was to investigate cerebral metabolism during the early and late stages of cerebral amyloid pathology in the APP / presenilin-1 (PS1) knock-in (APP/PS1 KI) mouse model of AD following a closed head injury (CHI). These mice develop both neuritic and non-neuritic amyloid plaque deposition starting at 6 months of age that increases linearly over time, accompanied by increased oxidative stress and metabolic disturbances as early as 1 to 2 months, with cognitive deficits manifesting at 11 months [33]. In this study, APP/PS1 KI mice 4-5 months of age (representing a stage just before the onset of amyloid pathology) were subjected to CHI and assessed at 1, 4 and 8-months post-injury. We hypothesized that TBI-induced mitochondrial dysfunction and Aβ-mediated impairments may interact through either distinct pathways that synergize or through overlapping pathways where effects might not be additive. Our findings reveal time-dependent interactions between injury and genotype, with pronounced mitochondrial dysfunction in injured KI mice at early time points, while amyloid-mediated effects predominate at later stages, providing insights into how these initially distinct insults may converge over time.

## Methods

### Animals

All experimental procedures involving animals were approved by the University of Kentucky’s Institutional Animal Care and Use Committee, following the guidelines set by the Guide for the Care and Use of Laboratory Animals and ARRIVE. Young adult (4-month-old) APP^NLh/NLh^ × PS1^P264L/P264L^ mice (RRID:MGI:3611210), initially developed by Cephalon [34], were used in this investigation. This mouse strain was crafted using gene-targeted knock-in (KI) technique to implant the Swedish FAD K670N/M671L mutations, convert the mouse β-amyloid sequence to human equivalent (NLh), and insert a proline to leucine (P264L) mutation into the mouse PS-1 gene [35, 36]. Mice were kept on a CD-1/129 genetic background (RRID:IMSR_CRL:022 x RRID:IMSR_JAX:002448). Wild-type (WT) mice were derived from heterozygous APP-PS1 mating pairs and preserved as a distinct lineage for over 20 inbreeding generations. The WT and APP/PS1 KI mice were backcrossed on average every 5 generations. Mice were grouped and kept under a 12/12 hour light/dark cycle with an unrestricted supply of conventional chow diet. The research was carried out with multiple batches of mice using a block design encompassing all experimental categories. Each cage housed several experimental groups, with every mouse receiving a unique identifier that masked its experimental group. Mice were randomly assigned to four experimental groups: WT+sham, WT+CHI, KI+sham, and KI+CHI, and were evaluated at 1, 4, and 8 months post-injury. At the time of surgery, mice were approximately 4.8 months old (group means ranging from 4.3 to 4.9 months, standard deviation (SD) 0.3–1.8 months). A total of 207 mice were included in the final analysis. One female APP/PS1 KI mouse died acutely during surgery due to anesthesia-related apnea and was excluded. Additionally, eight mice died from unknown causes during the study period: one KI+CHI mouse prior to 1 month, one KI+CHI and one WT+sham mouse prior to 4 months, and two WT+CHI, one WT+sham and KI+CHI mice prior to 8 months. No other animals were excluded due to surgical complications or mitochondrial isolation failure. However, four additional mice were excluded following data quality control due to excessive technical variability in mitochondrial respiration measurements (coefficient of variation >25%). Final sample sizes per group ranged from 9 to 18 animals per condition and time point, as reported in figure legends.

### Closed Head Injury (CHI)

As previous described [37–40], on the day of surgery the order in which mice underwent procedures was randomized with respect to genotype and assigned treatment (sham or CHI surgery), ensuring that mice from different experimental groups were interspersed throughout the surgical session rather than processed in blocks. This approach minimized potential time-of-day effects. All groups were included at the time of surgery, and the surgeon was blinded to the genotype of the mice. Mice were anesthetized using isoflurane (2.5-4%), and their heads were securely fastened in a stereotaxic apparatus. A midline sagittal cut was made before placing a 1 mL water-filled latex pipette bulb (ThermoFisher Scientific, Waltham, USA) beneath the mouse’s head. A stereotaxic electromagnetic impactor with a 5.0 mm flat steel end (Leica Biosystems, Wetzlar, Germany) was deployed to deliver a single controlled midline impact to the closed skull at coordinates: ML= 0.0 mm; AP= -1.6mm, with an impact depth (1.2 mm), a controlled speed (5.0 ± 0.2 m/sec), and dwell time (100 ms). Sham-operated mice underwent identical anesthesia and incision without impact. Mice were monitored during recovery and returned to group housing upon regaining full mobility.

### Mitochondria Isolation

Mitochondria were extracted using a multi-step method involving differential centrifugation and nitrogen disruption, as previously established [14, 41]. Animal euthanasia was achieved through carbon dioxide (CO_2_) administration, immediately followed by decapitation. Dissection and isolation of the hippocampus and neocortex were performed on a cold block, and tissues were homogenized in a cooled Dounce homogenizer filled with isolation buffer (215 mM mannitol, 75 mM sucrose, 0.1% bovine serum albumin (BSA), 1 mM ethylene glycol-bis(β-aminoethyl ether)-N,N,N′,N′-tetraacetic acid (EGTA), and 20 mM 4-(2-hydroxyethyl)-1-piperazineethanesulfonic acid (HEPES) adjusted to pH 7.2 with potassium hydroxide (KOH)). The homogenate was transferred to a 2 mL microcentrifuge tube and centrifuged at 1,300 × *g* for 3 minutes. The resulting supernatant was transferred to a new microcentrifuge tube and centrifuged again at 13,000 × *g* for 10 minutes. Following this, the supernatant was removed, and the resultant mitochondrial pellet was resuspended in 400 µL of isolation buffer. This suspension was then subjected to a nitrogen cell disruptor (Parr Instrument Company) pressurized at 1,200 pound-force per square inch (psi) for 10 minutes at 4°C, facilitating the release of synaptic mitochondria from synaptosomes. The total mitochondrial suspension (synaptic and non-synaptic mitochondria) was transferred to 1.5 mL centrifuge tubes and centrifuged at 13,000 × *g* for 10 minutes. The supernatants were discarded, and the resulting mitochondrial pellets were resuspended in 50–100 µL of EGTA-free isolation buffer to achieve a final concentration of ≥10 mg/mL. Mitochondrial protein concentration was estimated using a bicinchoninic acid (BCA) protein assay kit (Pierce, Rockford, IL).

### Mitochondrial respiration measurements

Mitochondrial bioenergetics were measured on isolated cortical and hippocampal mitochondria following established methods [42]. Oxygen consumption rates (OCRs) were measured using a Seahorse XFe96 Flux Analyzer (Agilent Technologies, Santa Clara, CA, USA), in the presence of various mitochondrial substrates, inhibitors, and uncouplers. All reagents were diluted in a respiration buffer containing 125 mM potassium chloride (KCl), 0.1% BSA, 20 mM HEPES, magnesium chloride (MgCl_2_), and 2.5 mM potassium dihydrogen phosphate (KH_2_PO_4_), adjusted to pH 7.2. Final concentrations were: 5 mM pyruvate, 2.5 mM malate, and 1 mM adenosine diphosphate (ADP) (Port A), 2.5 µM oligomycin A (Port B), 4 µM carbonyl cyanide-p-trifluoromethoxyphenylhydrazone (FCCP) (Port C), and 1 µM rotenone with 10 mM succinate (Port D). Each well was plated with 6 µg of total mitochondria in 30 µL, followed by centrifugation, then respiration buffer was gently added to reach 175 µL total volume. State III respiration (ADP-stimulated maximal respiration) was achieved with pyruvate, malate, and ADP (Port A), followed by State IV (basal respiration) with oligomycin (Port B). Uncoupled respiration representing State V Complex I was achieved with FCCP (Port C), and State V Complex II was achieved with rotenone and succinate (Port D). Each mouse was run in 4–8 technical replicates, and the average of these replicates was used for each animal.

### Histology and Immunohistochemistry

To confirm the presence and progression of amyloid pathology, an independent cohort of APP/PS1 KI mice was used to match the ages of animals at the three experimental post-injury time points: 5–6 months, 8–9 months, and 12–13 months of age. Mice were deeply anesthetized with 5% isoflurane in 100% oxygen and transcardially perfused with ice-cold 1X phosphate-buffered saline (PBS) for 5 minutes. Brains were rapidly extracted, and the left hemibrain was immersion-fixed in 4% paraformaldehyde (PFA) for 24 hours at 4°C, followed by cryoprotection in 30% sucrose until fully sunk. Tissue was coronally sectioned at 30 μm using a sliding microtome and stored at −20°C in cryoprotectant solution until staining. Immunohistochemical staining for amyloid-β was performed on free-floating sections using standard protocols. Sections were incubated with mouse biotinylated anti-β-amyloid 1–16 antibody (6E10; Covance/BioLegend, cat#39340-200, 1:3000 dilution, RRID:AB_662801). Antibody labeling was visualized using the avidin-biotin complex (ABC) method and 3,3′-diaminobenzidine (DAB) chromogen according to manufacturer instructions. Stained sections were scanned using a Zeiss Axio Scan Z.1 at 20× magnification and analyzed using HALO image analysis software (Indica Labs).

### Statistics

A median absolute deviation (MAD)-based outlier detection approach implemented in R (version 4.3.0, tidyverse 2.0.0) was used for quality control, removing outliers greater than three MAD units from the group median. After MAD-based quality control, samples with a coefficient of variation exceeding 25% across technical replicates were excluded from further analysis (n=4). Remaining replicate data were averaged to generate final subject-level measures. Batch effects introduced by assay day were corrected using ComBat (sva package, 3.48.0), stratified by brain region. For small batch sizes (<3 samples), mean-only correction was applied. The success of batch correction was evaluated using principal component analysis (PCA) and analysis of variance (ANOVA) to confirm elimination of batch effects without loss of biological variability. The effect of treatment groups was analyzed in JMP PRO (version 17.2.0) using two-way ANOVA (genotype x injury) separately for each brain region and time point. When a main effect of injury or a genotype x injury interaction was detected, post hoc t-tests were conducted to compare CHI versus sham within each genotype. This approach reflects our focus on injury effects rather than genotype effects; no post hoc tests were performed for main effects of genotype. Because these contrasts were hypothesis-driven and predefined in the analysis plan, adjustments such as Tukey’s test or false discovery rate (FDR) correction were not applied. As a secondary analysis to evaluate sex as a biological variable (SABV), we performed three-way ANOVAs (genotype × injury × sex) to test for sex interactions. Additionally, all data were stratified by sex and analyzed separately for males and females using two-way ANOVAs (genotype × injury) to ensure transparency in reporting potential sex-specific effects. All statistical tests used batch-corrected data, and significance was set at p < 0.05, and complete ANOVA results, including all F values, are provided in Supplementary Table 1.

## Results

Righting reflex time was recorded immediately following CHI as a measure of injury severity and acute neurological function. The latency to return to a prone position (righting reflex) was measured in seconds from the moment of impact until the animal successfully righted itself. Righting reflex latency was prolonged in CHI mice compared to sham controls. In WT+sham mice, females (F) righted in 87.8 ± 28.2 s and males (M) in 76.8 ± 19.2 s; following CHI, WT mice required 237.6 ± 94.6 s (F) and 272.3 ± 83.5 s (M). In APP/PS1 KI+sham mice, females righted in 77.1 ± 47.4 s and males in 122.4 ± 45.5 s; after CHI, KI mice required 297.1 ± 112.9 s (F) and 363.8 ± 158.1 s (M). One-way ANOVA demonstrated a significant effect of group on righting time (F= 48.15, p < 0.0001), and post hoc comparisons confirmed that both WT+CHI and KI+CHI mice took significantly longer to right than their respective sham controls (p < 0.0001), with APP/PS1 KI+CHI mice also slower than WT+CHI mice (p < 0.0001).

### Traumatic brain injury disrupts mitochondrial respiration in a brain region- and genotype-specific manner

We hypothesized that TBI would differentially affect mitochondrial function in wild-type and APP/PS1 KI mice across brain regions and over time. To test this, we measured mitochondrial respiratory function in the neocortex and hippocampus following CHI. These regions were selected for their distinct characteristics: the neocortex receives direct impact during CHI and exhibits early amyloid pathology, while the hippocampus experiences indirect force transmission and shows minimal Aβ plaque formation at 4-5 months of age when injury was delivered **(Fig. 1)**. In the neocortex **(Fig. 2A)**, we observed a trend toward a genotype-by-injury interaction at 1 month post-injury (F = 1.16, p = 0.069) with the APP/PS1 KI+CHI showing the lowest State III respiration (ADP-stimulated maximal oxygen consumption), with a 4.8% decrease in APP/PS1 KI+CHI mice (422.25 ± 10.61, least square mean (LSM) ± standard error of the mean (SEM)) compared to APP/PS1 KI+sham mice (442.96 ± 9.28, LSM ± SEM). Overall, the APP/PS1 KI mice showed reduced State III respiration compared to WT mice across all time points (1 month, F(1, 57) = 4.51, p = .039, partial η² = 0.083; 4 months, F(1, 57) = 10.35, p = .002, partial η² = 0.154; 8 months, F(1, 57) = 12.95, p < .001, partial η² = 0.245).

**Figure 1.**
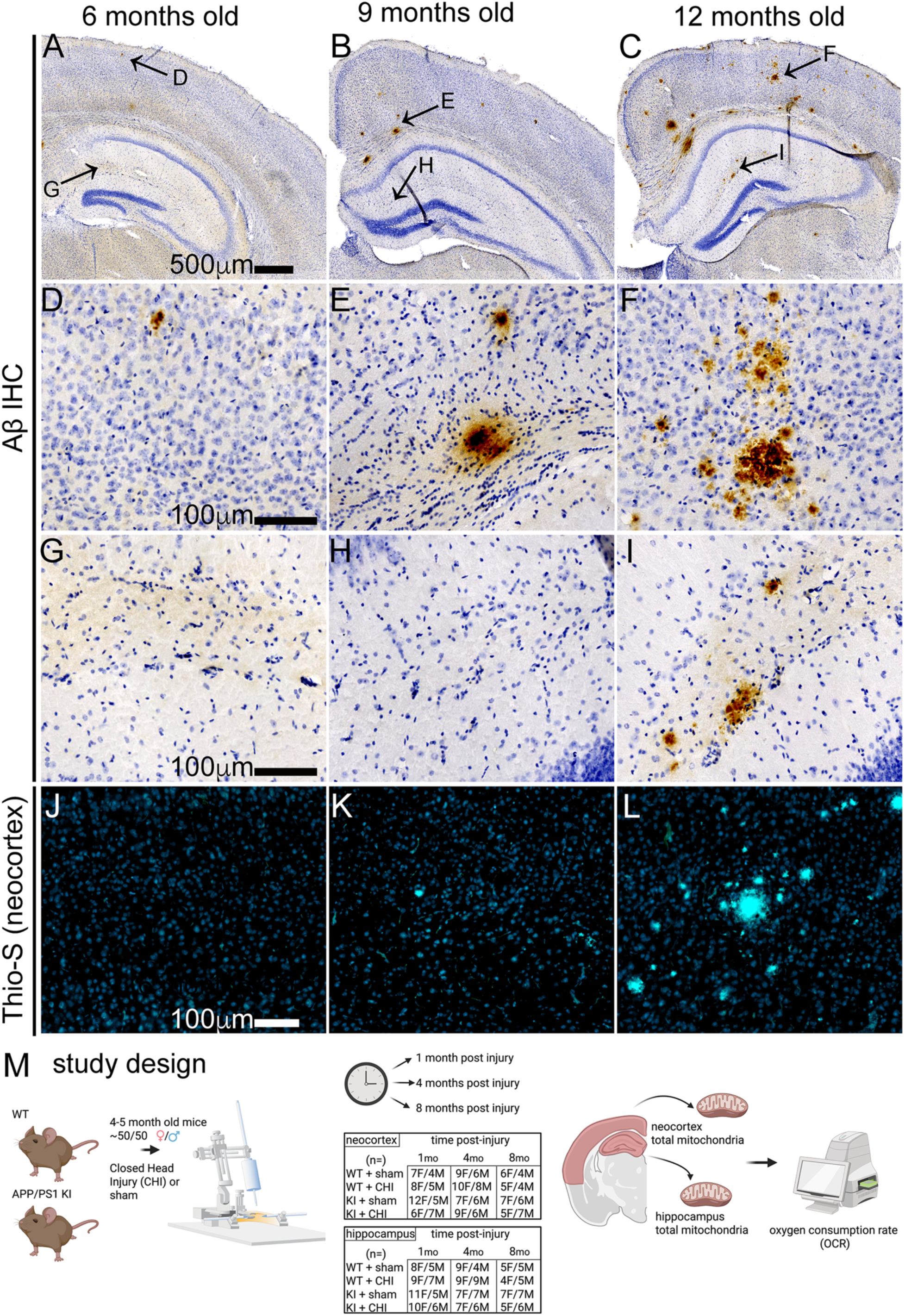
Association of amyloid-β accumulation in APP/PS1 KI mice as it influences the study design timeline. (A-C) Representative low magnification sections showing Aβ immunohistochemistry at 6, 9, and 12 months of age in male APP/PS1 KI mice. Arrows indicate locations of higher magnification images shown in the neocortex **(D-F)** and the hippocampus **(G-I)**. At 6 months, minimal Aβ pathology is observed. By 9 months, sparse plaques emerge, with substantial plaque burden developing by 12 months. **(J-L)** Thioflavin-S staining of neocortical regions confirms fibrillar amyloid deposits starting at 9 months, with a substantial burden by 12 months. **(M)** Study design showing APP/PS1 KI and wildtype (WT) mice receiving closed head injury (CHI) or sham surgery at ∼4-5 months of age, with assessments at 1, 4, and 8 months post-injury. The table shows final numbers of animals included in each group and time point. Endpoints included analysis of mitochondrial bioenergetics by Seahorse assay in the neocortex and hippocampus.

**Figure 2.**
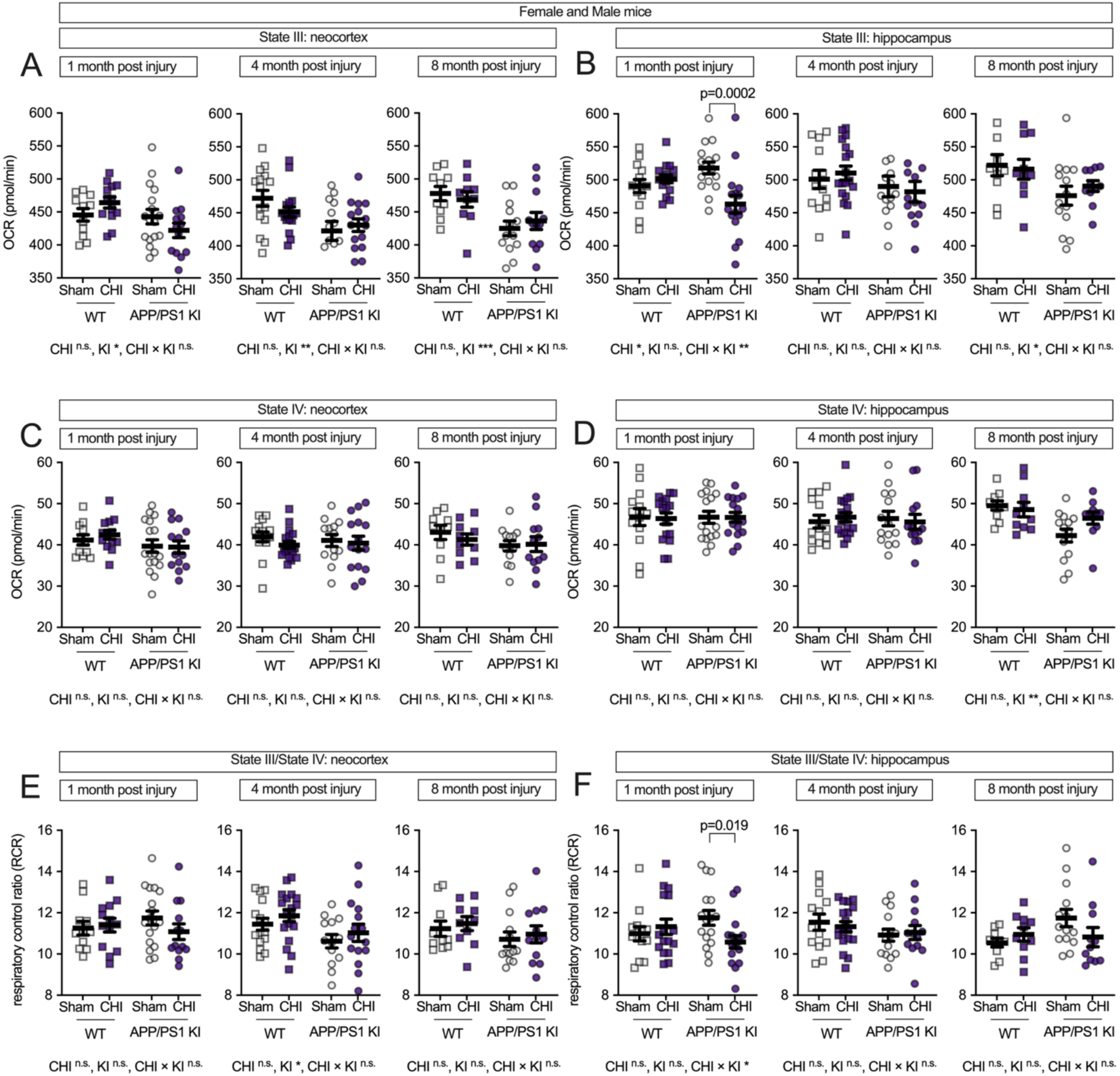
Interactive effects of genotype and CHI on mitochondrial respiratory capacity and coupling efficiency are observed in the hippocampus, but not in the neocortex. Mitochondrial oxygen consumption rate (OCR) was assessed in isolated mitochondria from WT and APP/PS1 KI mice at 1, 4, and 8 months post-injury. **(A)** In the neocortex, State III respiration showed genotype-dependent effects across all time points, with APP/PS1 KI mice exhibiting lower respiration. **(B)** In the hippocampus, CHI significantly reduced State III respiration at 1-month post-injury, specifically in APP/PS1 KI mice (p = 0.0002), with no significant injury effects at later time points. State IV respiration remained stable across all conditions in both the neocortex **(C)** and hippocampus **(D)**. The Respiratory control ratio (RCR, State III/State IV) in the neocortex **(E)** showed a genotype effect at 4 months post-injury. In the hippocampus **(F)**, RCR was significantly reduced in APP/PS1 KI mice with CHI compared to sham controls at 1-month post-injury. Sample sizes as follows: 1 month—WT+sham (n=11), WT+CHI (n=13), KI+sham (n=17), KI+CHI (n=13); 4 months—WT+sham (n=15), WT+CHI (n=18), KI+sham (n=13), KI+CHI (n=15); 8 months—WT+sham (n=10), WT+CHI (n=9), KI+sham (n=13), KI+CHI (n=12). Summary statistics show mean ± SEM, with individual data points representing each animal (squares = WT; circles = APP/PS1 KI; open symbols = sham; filled symbols = CHI). Statistics shown in the figure represent planned post hoc t-tests comparing CHI vs. sham within genotypes.

State III respiration in the hippocampus **(Fig. 2B)** showed a different pattern, with genotype differences appearing later and becoming detectable at 8 months post-injury (F(1, 57) = 6.38, p = 0.016, partial η² = 0.138). Notably, at 1 month post-injury, we observed both a main effect of injury (F(1, 57) = 4.92, p = 0.030, partial η² = 0.080) and a significant injury-by-genotype interaction (F(1, 57) = 10.98, p = 0.002, partial η² = 0.161). APP/PS1 KI mice with CHI showed a 10.6% reduction in State III respiration compared to APP/PS1 KI+sham controls (p = 0.0002), reflecting reduced mitochondrial oxygen consumption (APP/PS1 KI+CHI: 463.3 ± 53.1 vs. APP/PS1 KI+sham: 518.0 ± 35.4, LSM ± SEM). However, no significant injury or interaction effects were detected at 4 or 8 months post-injury.

State IV respiration (basal oxygen consumption without ADP) remained unchanged across all groups in both brain regions **(Fig. 2C-D)**, indicating that mitochondrial dysfunction primarily affects stimulated rather than basal respiratory capacity. The respiratory control ratio (RCR, State III/State IV), which reflects mitochondrial coupling efficiency, showed a genotype effect in the neocortex at 4 months post-injury (F(1, 57) = 6.00, p = 0.017, partial η² = 0.095; **Fig. 2E**). In the hippocampus at 1 month post-injury, a significant genotype-by-injury interaction (F(1, 57) = 10.98, p = 0.002, partial η² = 0.161) revealed reduced RCR in injured APP/PS1 KI mice compared to sham controls (p = 0.019; **Fig. 2F**), suggesting that hippocampal mitochondrial coupling efficiency is particularly vulnerable to CHI in the context of APP/PS1 mutations.

### Electron transport chain dysfunction is exacerbated by TBI in APP/PS1 KI mice

To further understand the impact on specific components of the ETC, we measured State V respiration for individual complexes. In the neocortex (**Fig. 3A**), Complex I-driven respiration (NADH-linked oxidative phosphorylation) showed reduced function in APP/PS1 KI mice across all time points (1 month, F(1, 57) = 5.06, p = .029, partial η² = 0.092; 4 months, F(1, 57) = 23.88, p < .001, partial η² = 0.295; and 8 months, F(1, 57) = 21.07, p < 0.001, partial η² = 0.345.) compared to WT mice. At 1 month post-injury, in the neocortex, the genotype-by-injury interaction did not reach statistical significance (F(1, 57) = 3.60, p = 0.063, partial η² = 0.067, medium effect size). In the hippocampus (Fig. 3B), there was a significant genotype-by-injury interaction (F(1, 57) = 6.10, p = 0.017, partial η² = 0.097), with the APP/PS1 KI mice with CHI showing reduced Complex I-driven respiration compared to APP/PS1 KI+sham controls (p = 0.004).

**Figure 3.**
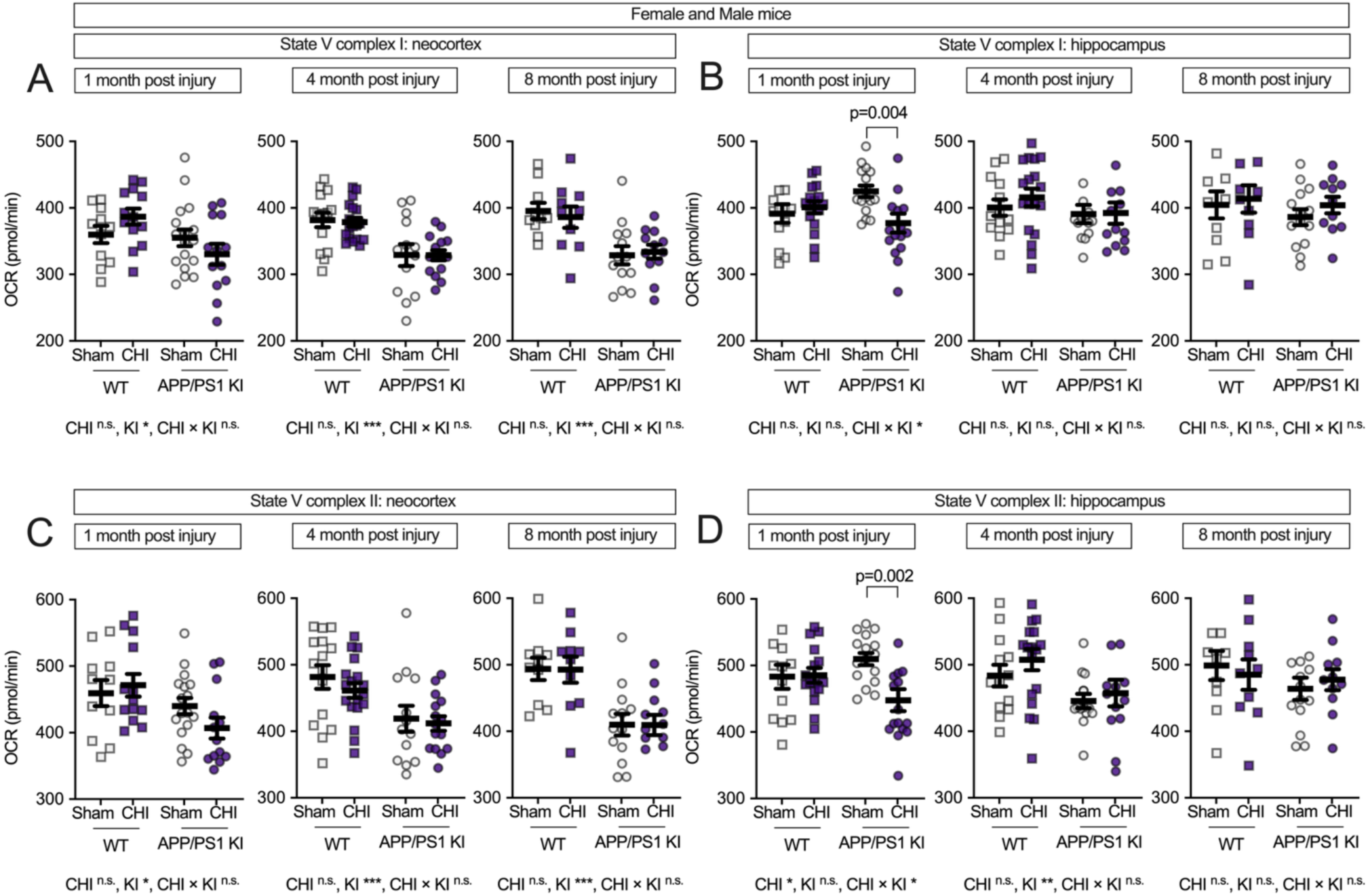
CHI disrupts electron transport chain function in a region-dependent manner. State V respiration was measured to assess mitochondrial electron chain complex-specific function in isolated mitochondria from WT and APP/PS1 KI mice at 1, 4, and 8 months post-injury. Complex I-driven respiration (glutamate/malate substrate) was impaired in APP/PS1 KI mice across all time points in the neocortex (A), with no injury effects. In contrast, (B) in the hippocampus, CHI significantly reduced Complex I respiration at 1 month post-injury in APP/PS1 KI mice. (C) The APP/PS1 KI mice had lower Complex II-driven respiration (succinate substrate) across time in the neocortex. (D) The CHI reducing Complex II activity at 1 month post-injury in the hippocampus. Sample sizes per group ranged from 9 to 18 per time point, as detailed in Figure 2. Summary statistics show mean ± SEM, with individual data points representing each animal. Statistics shown on the figure represent planned post hoc t-tests comparing CHI vs. sham within genotypes.

Complex II-driven respiration (FAD-linked oxidative phosphorylation) showed genotype-associated deficits in both brain regions. In the neocortex (**Fig. 3C**), APP/PS1 KI mice showed lower respiration than WT mice across all time points (1 month: F(1, 57) = 6.98, p = 0.011, partial η² = 0.123; 4 months: F(1, 57) = 14.32, p = < 0.001, partial η² = 0.201; 8 months: F(1, 57) = 25.28, p = < 0.001, partial η² = 0.387). In the hippocampus (**Fig. 3D**), we observed a genotype-by-injury interaction at 1 month post-injury (F(1, 57) = 5.30, p = 0.025, partial η² = 0.085), with APP/PS1 KI mice with CHI showing lower respiration than sham controls (p = 0.002), while genotype differences became detectable at 4 months post-injury (F(1, 57) = 7.47, p = 0.008, partial η² = 0.122).

### Female-specific vulnerability to traumatic brain injury in APP/PS1 KI mice

We next examined potential sex differences in mitochondrial function following CHI, as previous studies have reported sexually dimorphic effects in both brain injury recovery and amyloid-related pathology [43–51]. While we did not detect significant main effects of sex alone (all F < 1.86, all p > 0.17), a three-way ANOVA revealed significant interactions between sex × injury and sex × genotype. Specifically, we observed significant genotype × sex interactions for multiple mitochondrial endpoints (F values: 4.74-12.00, p = 0.001-0.034) and sex × injury type interactions (F = 5.99-6.18, p = 0.016-0.018), with the most robust effects observed in hippocampal tissue at both 1 and 8 months post-injury. Therefore, to assess these sex-dependent patterns, we stratified the data by sex and analyzed males and females separately. In female mice, the neocortex **(Fig. 4A)** showed a genotype effect at all post-injury time points, with APP/PS1 KI mice exhibiting lower State III respiration than WT controls (1 month: F(1, 29) = 6.96, p = 0.013, partial η² = 0.193; 4 months: F(1, 31) = 5.77, p = 0.022, partial η² = 0.157; 8 months: F(1, 19) = 4.99, p = 0.038, partial η² = 0.208). At 4 months post-injury, we observed a genotype-by-injury interaction (F(1, 31) = 4.51, p = 0.042, partial η² = 0.127), with WT+CHI mice showing reduced respiration compared to WT+sham controls (p = 0.009). In the hippocampus of female mice (**Fig. 4B**), the data visually suggest an interaction between injury and genotype, with KI+CHI mice showing lower State III respiration than KI+sham controls. The interaction had a medium effect size (partial η² = 0.094) but did not reach statistical significance at 1 month post-injury (F(1, 34) = 3.52, p = 0.069). By 8 months post-injury, a genotype effect emerged (F(1, 17) = 16.09, p = < 0.001, partial η² = 0.486), with APP/PS1 KI mice exhibiting a 13.5% reduction in State III respiration compared to WT (APP/PS1 KI = 466.34 ± 11.98, WT = 539.38 ± 13.72, LSM ± SEM).

**Figure 4.**
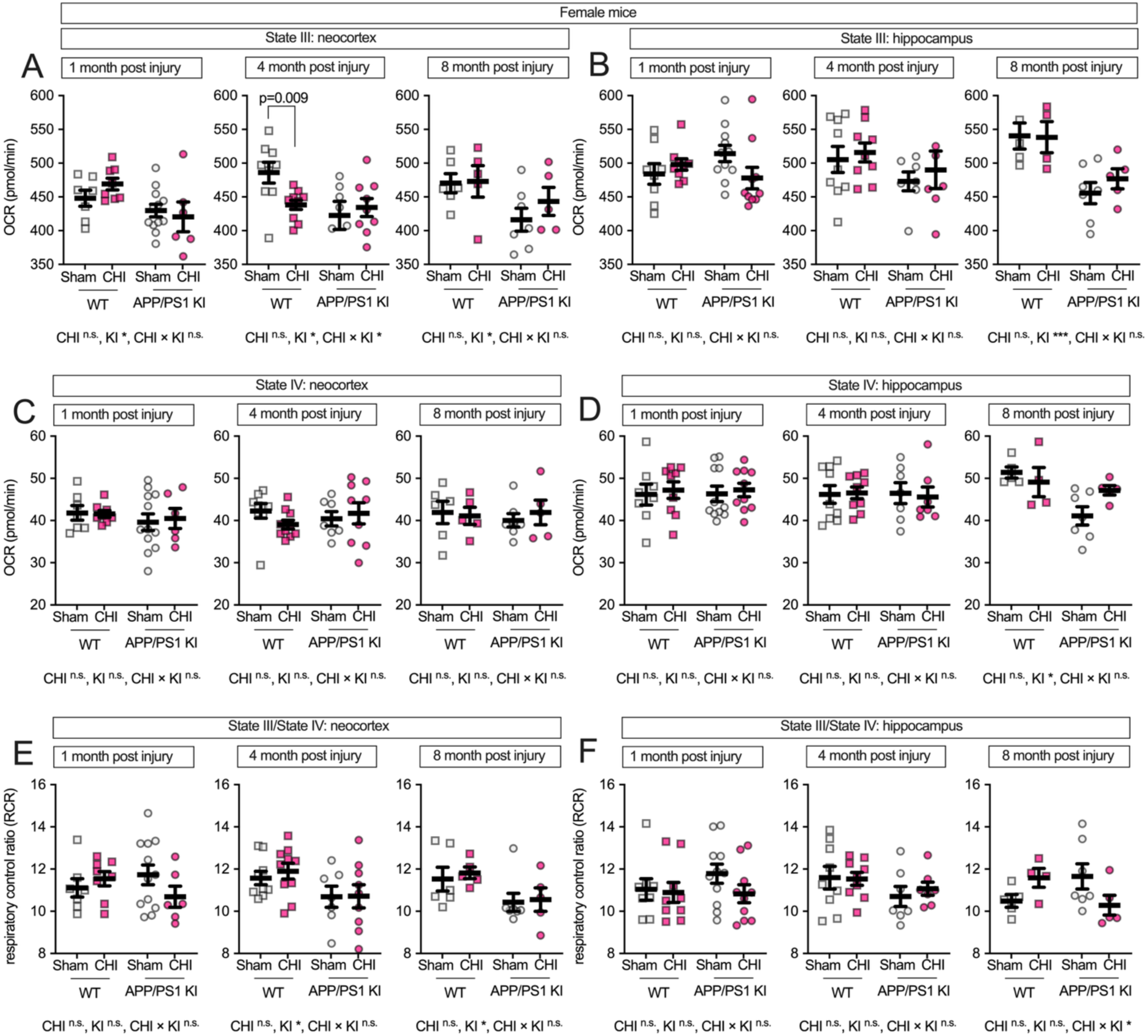
Maximum respiratory capacity and coupling efficiency in female mice following CHI. State III respiration in isolated mitochondria from female mice showed genotype effects in the neocortex **(A)** at all time points, with a genotype-by-injury interaction at 4 months post-injury. In the hippocampus **(B)**, a trend toward genotype-by-injury interaction was observed at 1 month, with a significant genotype effect at 8 months. State IV respiration **(C-D)** remained stable until 8 months, when genotype effects emerged in hippocampus. RCR showed genotype effects in neocortex at 4 and 8 months **(E)** and a genotype-by-injury interaction in hippocampus at 8 months **(F)**. Sample sizes: 1 month—WT+sham (n=7), WT+CHI (n=8), KI+sham (n=12), KI+CHI (n=6); 4 months—WT+sham (n=9), WT+CHI (n=10), KI+sham (n=7), KI+CHI (n=9); 8 months—WT+sham (n=6), WT+CHI (n=5), KI+sham (n=7), KI+CHI (n=5). Summary statistics show mean ± SEM, with individual data points representing each animal (squares = WT; circles = APP/PS1 KI; open symbols = sham; filled symbols = CHI). Summary statistics show mean ± SEM, with individual data points representing each animal. Statistics shown on the figure represent planned post hoc t-tests comparing CHI vs. sham within genotypes.

State IV respiration in female mice showed no differences at 1 and 4 months post-injury. At 8 months in the hippocampus, we observed a genotype effect (F(1, 17) = 8.07, p = 0.011, partial η² = 0.322; **Fig. 4C-D**). RCR in the neocortex showed genotype effects at 4 and 8 months (F(1, 31) = 5.52, p = 0.025, partial η² = 0.151; F(1, 19) = 5.95, p = 0.025, partial η² = 0.238; **Fig. 4E**), while the hippocampus exhibited a genotype-by-injury interaction at 8 months (F(1, 17) = 3.79, p = 0.068, partial η² = 0.182; **Fig. 4F**), with APP/PS1 KI+CHI mice showing an 11.7% lower RCR than sham controls (LSM ± SEM: KI+CHI = 10.29 ± 0.53 vs. KI+sham = 11.65 ± 0.44).

In female mice, Complex I-driven respiration in the neocortex (**Fig. 5A**) visually suggests an interaction between genotype and injury at 1 month post-injury, with KI+CHI mice showing an 11.7% reduction compared to KI+sham controls (KI+CHI = 311.61 ± 18.51 vs. KI+sham = 352.93 ± 13.09, LSM ± SEM). The interaction had a medium effect size (partial η² = 0.100) but did not reach statistical significance (F(1, 29) = 3.23, p = 0.083). A significant genotype effect was present at this time point (F(1, 29) = 8.73, p = 0.006, partial η² = 0.231) and persisted at 4 months (F(1, 31) = 14.87, p < 0.001, partial η² = 0.324) and 8 months (F(1, 19) = 13.15, p = 0.002, partial η² = 0.409). In the hippocampus **(Fig. 5B)**, a genotype effect emerged at 8 months (F(1, 17) = 7.59, p = 0.014, partial η² = 0.309). Complex II-driven respiration in female mice showed genotype effects in the neocortex **(Fig. 5C)** across all time points (1 month: F(1, 29) = 9.53, p = 0.004, partial η² = 0.247; 4 months: F(1, 31) = 5.77, p = 0.022, partial η² = 0.157; 8 months: F(1, 19) = 11.57, p = 0.003, partial η² = 0.379), with APP/PS1 KI mice showing a 13.0% reduction at 1 month and a 16.9% reduction by 8 months compared to WT. In the hippocampus **(Fig. 5D)**, the pattern reversed with APP/PS1 KI mice initially showing slightly higher activity at 1 month (4.0%), but a 16.8% reduction by 8 months compared to WT.

**Figure 5.**
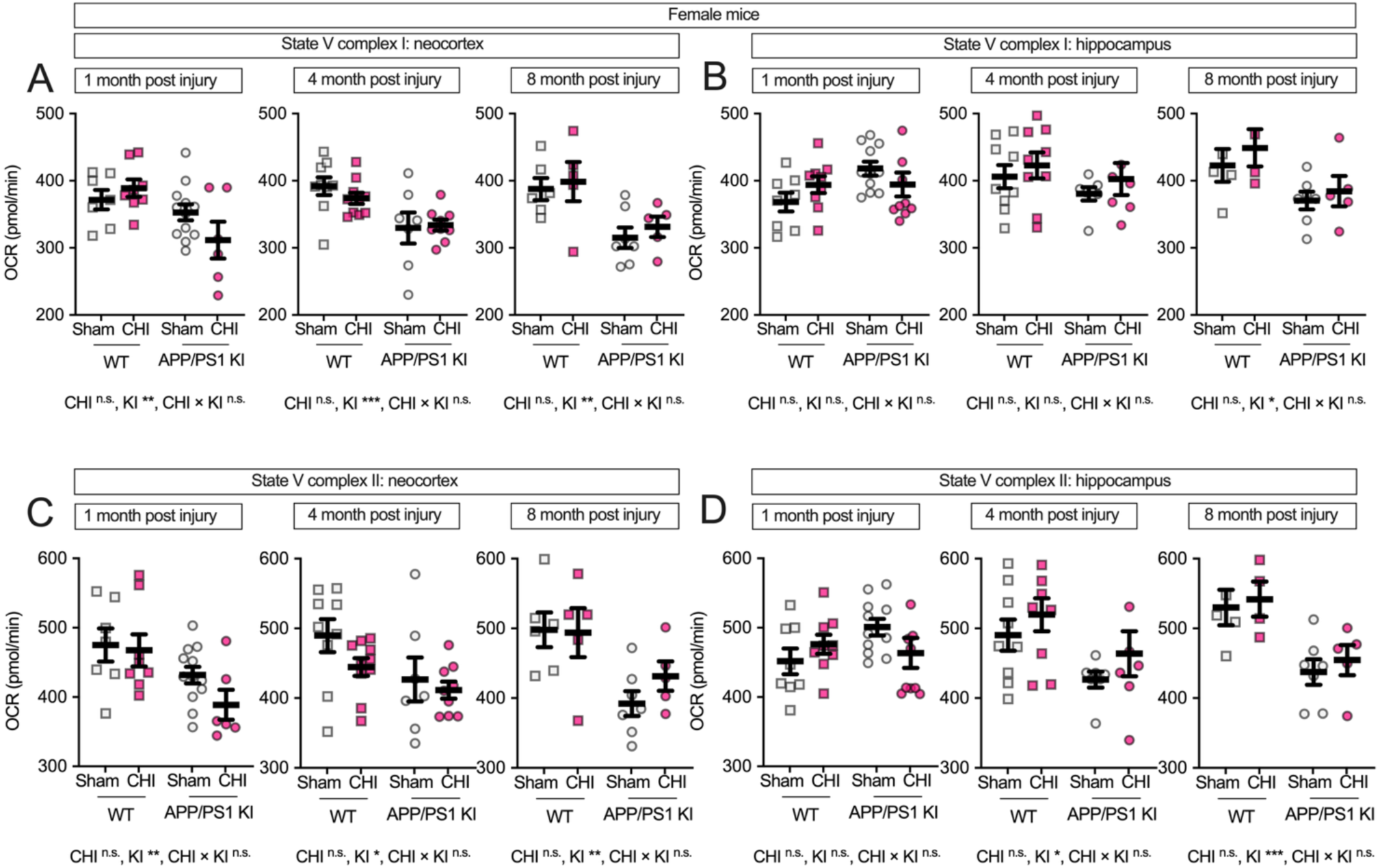
Genotype but not CHI disrupts electron transport chain function in female APP/PS1 KI mice. State V respiration was assessed in mitochondria from female WT and APP/PS1 KI mice at 1, 4, and 8 months post-injury (average age: 8.47 ± 3.48 months). Complex I-driven respiration declined over time in APP/PS1 KI mice in the neocortex (A), but CHI had no additional effect. In the hippocampus (B), CHI caused a transient decrease in Complex I-driven respiration at 1 month post-injury in APP/PS1 KI mice. Complex II-driven respiration followed a similar pattern, with APP/PS1 KI mice exhibiting deficits over time in the neocortex (C), while CHI selectively reduced respiration at 1 month post-injury in the hippocampus (D). All measurements were performed under FCCP-uncoupled conditions to assess maximal electron transport chain function. Sample sizes for females ranged from 4 to 12 animals per group per time point. Summary statistics show mean ± SEM, with individual data points representing each animal. Sample sizes for females ranged from 4 to 12 per group per time point, as follows: 1 month—WT+sham (n=7), WT+CHI (n=8), KI+sham (n=12), KI+CHI (n=6); 4 months—WT+sham (n=9), WT+CHI (n=10), KI+sham (n=7), KI+CHI (n=9); 8 months—WT+sham (n=6), WT+CHI (n=5), KI+sham (n=7), KI+CHI (n=5). Summary statistics show mean ± SEM, with individual data points representing each animal (squares = WT; circles = APP/PS1 KI; open symbols = sham; filled pink symbols = CHI). Statistics shown on the figure represent planned post hoc t-tests comparing CHI vs. sham within genotypes.

### Traumatic brain injury-induced hippocampal mitochondrial deficits in male APP/PS1 KI mice

Male mice exhibited distinct mitochondrial responses compared to females following CHI. In the neocortex (**Fig. 6A**), APP/PS1 KI+CHI mice exhibited a 2.1% reduction in State III respiration compared to KI+sham controls at 1 month post-injury (KI+CHI = 420.69 ± 14.33 vs. KI+sham = 429.82 ± 10.13, LSM ± SEM). The genotype-by-injury interaction showed a large effect size (partial η² = 0.153) but did not reach statistical significance (F(1, 17) = 3.07, p = 0.098). At 4 months post-injury, a genotype effect emerged (F(1, 22) = 4.33, p = 0.049, partial η² = 0.164), and by 8 months, this effect strengthened (F(1, 17) = 7.38, p = 0.015, partial η² = 0.303), with APP/PS1 KI mice showing an 8.8% reduction compared to WT controls (KI = 429.87 ± 13.04 vs. WT = 471.77 ± 13.48, LSM ± SEM). Strikingly, the hippocampus of male mice (**Fig. 6B**) showed a strong genotype-by-injury interaction at 1 month post-injury (F(1, 19) = 11.45, p = 0.003, partial η² = 0.376), with APP/PS1 KI+CHI mice showing a significant 16.6% reduction in State III respiration compared to KI+sham controls (p = 0.0003; KI+CHI = 438.71 ± 13.22 vs. KI+sham = 526.38 ± 14.48, LSM ± SEM). State IV respiration remained stable across all groups **(Fig. 6C-D)**. RCR in the neocortex showed no significant changes (**Fig. 6E**), while the hippocampus showed a genotype-by-injury interaction (F(1, 19) = 5.25, p = 0.034, partial η² = 0.216; **Fig. 6F**), with APP/PS1 KI+CHI mice exhibiting reduced coupling efficiency at 1 month post-injury.

**Figure 6.**
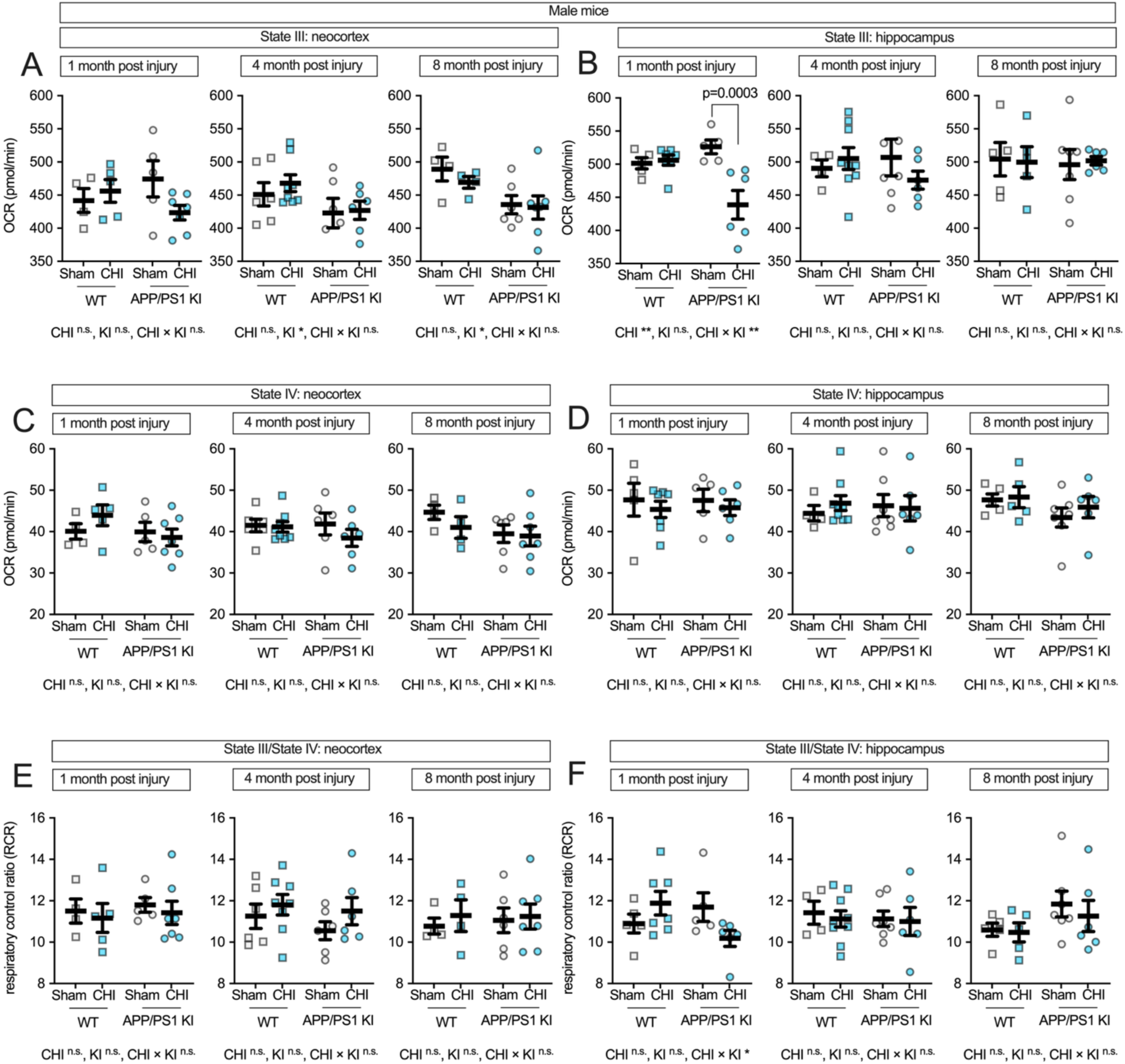
Maximum respiratory capacity and coupling efficiency in male mice following CHI. State III respiration in isolated mitochondria from male mice showed mild genotype effects in the neocortex **(A)**, while the hippocampus **(B)** exhibited a strong genotype-by-injury interaction at 1 month post-injury. State IV respiration **(C-D)** remained stable across all conditions. RCR showed no significant changes in neocortex **(E)** but demonstrated a genotype-by-injury interaction in hippocampus at 1 month post-injury **(F)**. Sample sizes: 1 month—WT+sham (n=4), WT+CHI (n=5), KI+sham (n=5), KI+CHI (n=7); 4 months—WT+sham (n=6), WT+CHI (n=8), KI+sham (n=6), KI+CHI (n=6); 8 months— WT+sham (n=4), WT+CHI (n=4), KI+sham (n=6), KI+CHI (n=7). Summary statistics show mean ± SEM, with individual data points representing each animal (squares = WT; circles = APP/PS1 KI; open symbols = sham; filled symbols = CHI).

To examine complex-specific mitochondrial function in male mice, we measured State V respiration for individual ETC complexes. In the neocortex, Complex I-driven respiration (**Fig. 7A)** showed genotype effects at 4 and 8 months post-injury (4 months: F(1, 22) = 8.24, p = 0.009, partial η² = 0.272; 8 months: F(1, 17) = 7.22, p = 0.016, partial η² = 0.298), with no significant injury effects or interactions at any time point. The hippocampus (**Fig. 7B**), however, showed vulnerability in male APP/PS1 KI mice 1 month post-injury, with a main effect of injury (F(1, 19) = 9.98, p = 0.005, partial η² = 0.344) and a genotype-by-injury interaction (F(1, 19) = 4.39, p = 0.050, partial η² = 0.188). Male APP/PS1 KI+CHI mice had a 9.5% reduction in Complex I-driven respiration compared to KI+sham controls (p = 0.0017; KI+CHI = 379.61 ± 11.52 vs. KI+sham = 419.63 ± 12.12, LSM ± SEM). This contrasts with female APP/PS1 KI mice, which showed only a 1.7% decrease in hippocampal Complex I-driven respiration following CHI (KI+CHI = 394.20 ± 9.82 vs. KI+sham = 406.26 ± 9.33), a 5.5-fold smaller deficit than in males.

**Figure 7.**
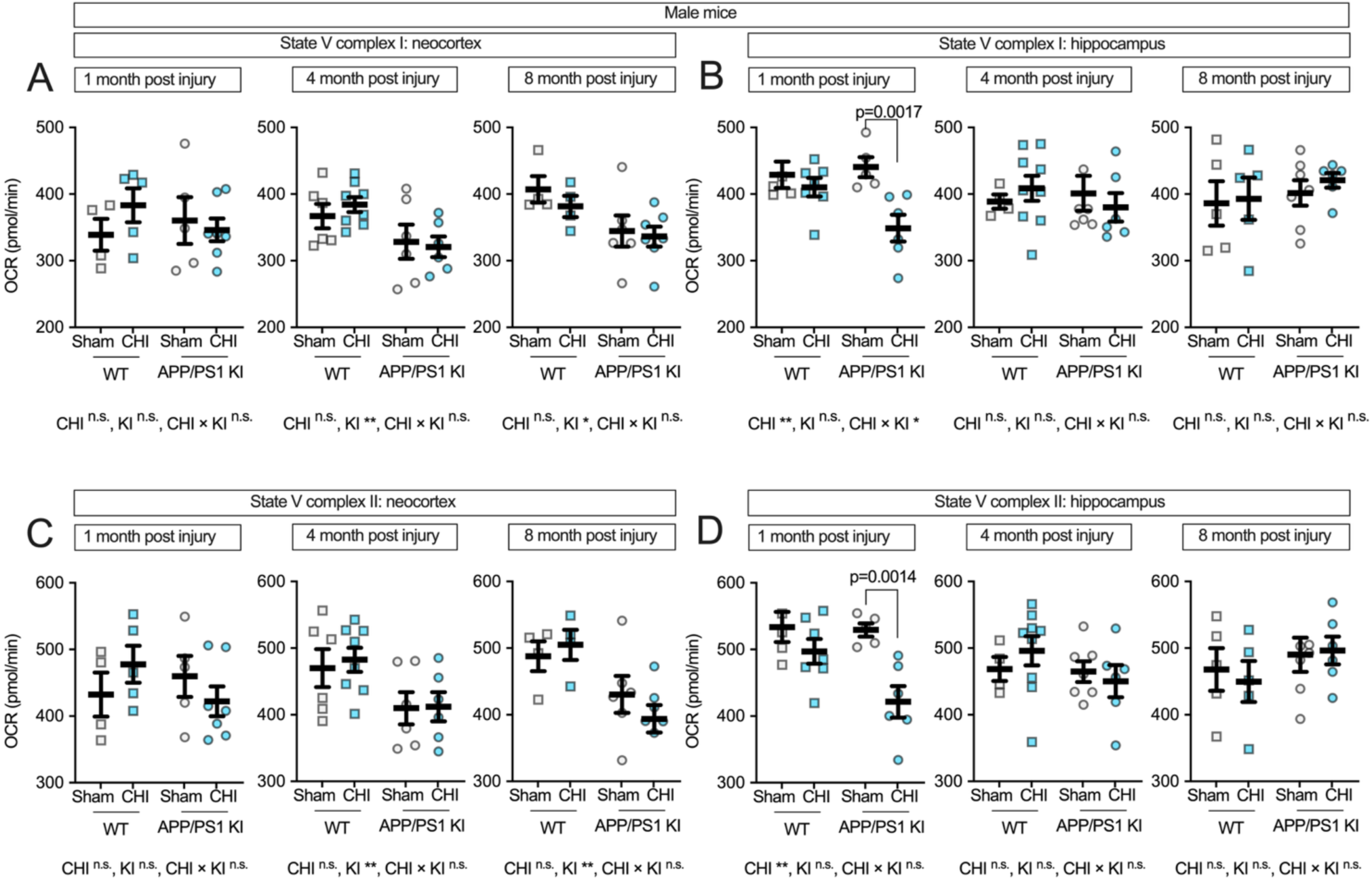
Complex-specific electron transport chain function in male mice following CHI. State V respiration in isolated mitochondria from male mice showed genotype effects for Complex I-driven respiration in the neocortex **(A)**, while the hippocampus **(B)** revealed both injury effects and genotype-by-injury interactions at 1-month post-injury. Complex II-driven respiration showed similar patterns, with genotype effects in the neocortex **(C)** and injury-induced reductions in the hippocampus **(D)** at 1-month post-injury in APP/PS1 KI mice. Sample sizes: 1 month—WT+sham (n=4), WT+CHI (n=5), KI+sham (n=5), KI+CHI (n=7); 4 months—WT+sham (n=6), WT+CHI (n=8), KI+sham (n=6), KI+CHI (n=6); 8 months—WT+sham (n=4), WT+CHI (n=4), KI+sham (n=6), KI+CHI (n=7). Summary statistics show mean ± SEM, with individual data points representing each animal (squares = WT; circles = APP/PS1 KI; open symbols = sham; filled symbols = CHI).

Complex II-driven respiration in males showed a similar pattern to Complex I. In the neocortex (**Fig. 7C**), genotype effects were observed at 4 and 8 months post-injury (4 months: F(1, 22) = 8.05, p = 0.010, partial η² = 0.268; 8 months: F(1, 17) = 11.21, p = 0.004, partial η² = 0.397) without significant injury effects. In the hippocampus (**Fig. 7D**), we observed a main effect of injury at 1 month post-injury (F(1, 19) = 12.91, p = 0.002, partial η² = 0.405) and a genotype-by-injury interaction of medium effect size (partial η² = 0.141) that did not reach statistical significance (F(1, 19) = 3.12, p = 0.094). Male APP/PS1 KI+CHI mice showed a 20.4% reduction in Complex II-driven respiration compared to KI+sham controls (p = 0.0014; KI+CHI = 450.27 ± 22.28 vs. KI+sham = 529.43 ± 21.35, LSM ± SEM). At later time points, these injury effects diminished as genotype-associated decline progressed, with injured and non-injured KI mice showing similar levels of impairment by 8 months post-injury.

## Discussion

The mechanisms by which TBI increases risk for AD remain incompletely defined. Our mitochondrial bioenergetic analysis of APP/PS1 KI mice following a single CHI revealed significant deficits in mice with both genetic susceptibility and injury that were not observed in other groups. These deficits showed distinct temporal patterns, with the most pronounced genotype-by-injury interactions appearing at early time points, particularly in the hippocampus. The differential vulnerability between brain regions likely reflects both the biomechanics of injury and the spatial pattern of amyloid pathology. Furthermore, biological sex emerged as a critical determinant of vulnerability, with male mice exhibiting substantially greater mitochondrial dysfunction following injury than females. Persistent mitochondrial impairment is increasingly recognized as both a marker and driving force in neurodegeneration, where chronic deficits in oxidative phosphorylation and the resulting oxidative stress may initiate a feed-forward cycle. In this cycle, energy failure promotes protein aggregation and inflammation, which further compromises mitochondrial function. This dynamic interplay between acute injury and chronic pathology may help explain why epidemiological studies in humans consistently identify TBI as a risk factor for dementia.

The temporal patterns we observed, with early injury-induced deficits giving way to genotype-dominated effects at later time points, suggest that initial TBI-associated mitochondrial dysfunction may accelerate or exacerbate processes already compromised by amyloid pathology. For example, within 24 hours of mild TBI in mice, reductions in mitochondrial membrane potential are accompanied by increased calcium retention and upregulation of NCLX (Na/Li/Ca exchanger) to mitigate calcium overload [52], while Aβ concurrently impairs calcium uptake, compromising buffering capacity, as shown in vitro with oligomeric Aβ1–42 and in transgenic amyloidosis mouse models [53–55]. The pronounced deficits we observed in Complexes I and II function, particularly in male mice, align with these calcium dysregulation mechanisms. Similarly, TBI initiates the removal of damaged mitochondria via mitophagy [56–58], but Aβ interferes with PINK1 stabilization, leading to inadequate clearance of defective organelles as observed in APPSwInd (J-20) mice in Du et al., 2017 and also reported in human samples, APP/PS1 transgenic mice, and C. elegans [59, 60]. Moreover, TBI disrupts the balance of mitochondrial dynamics by increasing fission, as evidenced by elevated Drp1 and Fis1 levels and reduced mitochondrial length [61, 62], a process further exacerbated by Aβ’s colocalization with Drp1, demonstrated in human brain tissue and in primary hippocampal neurons from AβPP×PS1 transgenic mice [63]. This shift toward excessive fission is compounded by decreased levels of fusion proteins (Mfn1/2 and OPA1) in both TBI and AD [59, 64, 65], promoting fragmentation, reduced ATP production, and heightened apoptotic vulnerability. Our observation that the hippocampus shows greater vulnerability than the neocortex may reflect the pattern of amyloid pathology, with genotype effects being more pronounced in the hippocampus. This suggests that when genotype effects are stronger, they may act on the same pathways as TBI-induced damage, thus masking or preventing additional injury effects from manifesting. Collectively, these findings are consistent with the hypothesis that TBI-induced compensatory mechanisms may initially clear damaged mitochondria, but that the presence of Aβ impairs these processes, ultimately driving progressive mitochondrial dysfunction that converges with amyloid-mediated pathology over time. However, we cannot exclude the possibility that TBI does not engage compensatory clearance in this context and that the absence of additive TBI effects beyond amyloid pathology in most comparisons reflects a lack of such a mechanism. Distinguishing between these possibilities will require future studies designed to directly test mitochondrial turnover and clearance.

In addition to mitochondrial quality control, both TBI and Aβ directly impact electron transport chain function. These deficits likely stem from multiple mechanisms: Complex I is highly susceptible to oxidative injury and electron leakage, with metabolic damage impairing coenzyme Q10 reduction and ATP synthesis [66–69]. While Aβ specifically inhibits Complex I and exacerbates reactive oxygen species (ROS) production [70], accounting for the robust genotype effects that worsen with increasing Aβ load, TBI-induced oxidative stress, calcium overload, and lipid peroxidation [71] likely contribute to the Complex I respiration decreases we observed. Both insults also impact metabolic enzymes feeding into the ETC. For example, pyruvate dehydrogenase (PDH) is inhibited in both TBI and AD [72, 73], while Aβ accumulating within mitochondria binds to PDH and α-ketoglutarate dehydrogenase, reducing TCA cycle flux and electron donor availability [74]. Additionally, both conditions impair mitochondrial membrane potential [71, 75, 76], which drives Complex V activity, thereby reducing ATP production and increasing ROS [77]. The lipid peroxidation product 4-Hydroxynonenal, elevated in both TBI and AD, modifies Complex V in AD brains and accumulates rapidly post-injury [78–80].

In WT (CD1/129) mice, no significant mitochondrial deficits were detected at the 1-month time point, whereas in our previous studies of B6 WT mice, we observed deficits at 1 month but not at 3 days post-injury, suggesting a delayed onset of dysfunction in the current strain [37]. These findings highlight the importance of genetic background in shaping the temporal trajectory of injury responses and suggest that future studies should examine earlier post-injury time points in CD1/129 mice and directly compare multiple strains to determine whether the onset and pattern of mitochondrial deficits after CHI are strain-dependent. In the present study, by 4 months post-injury, however, mitochondrial changes emerged in the neocortex of female WT mice, which were absent at both 1- and 8-month assessments. Sex differences significantly influence mitochondrial responses to neurodegeneration and injury, with AD models often showing more severe genotype-driven mitochondrial impairments in females, likely due to interactions between Aβ, estrogen, and mitochondrial pathways [81–84]. In contrast, TBI models frequently exhibit greater acute mitochondrial dysfunction in males, particularly in the hippocampus. At the same time, females may benefit from estrogen and progesterone’s neuroprotective effects, including enhanced antioxidant defenses and mitochondrial function [85–88]. We found the strongest interaction of TBI and Aβ in male mice, suggesting that female mice may show some degree of protection against mitochondrial changes at the 1 month post-injury time point, which we hypothesize could be mediated by hormonal mechanisms. However, at 4 months post-injury, we observed the emergence of mitochondrial deficits in WT female mice, which we hypothesize could be related to perimenopausal-like endocrine changes. These changes typically begin around 9 months of age in mice, though some strains may exhibit irregular cycling as early as 8 months [89–91]. Without assessing reproductive senescence, we cannot determine whether these effects are directly linked to specific stages of endocrine aging in our cohort, and this remains an important area for future study.

## Conclusions

This study demonstrates that TBI exacerbates mitochondrial dysfunction in APP/PS1 KI mice in a region- and sex-specific manner, with males showing greater vulnerability to injury-induced deficits in the hippocampus. Our temporal analysis revealed that early injury-dominated effects gradually converge with genotype-driven changes, providing a framework for understanding how TBI may increase AD risk. Future studies should build upon these findings by examining specific mechanisms underlying this vulnerability, particularly mitochondrial quality control pathways (mitophagy, fusion/fission dynamics, and biogenesis) that may be dysregulated following injury. Therapeutic strategies targeting mitochondrial function, including PDH kinase inhibitors [92] or alternative biofuel substrates like beta-hydroxybutyrate that bypass damaged metabolic enzymes [93], warrant investigation not only for their potential to rescue bioenergetic deficits but also for their ability to mitigate cognitive decline. By targeting these mitochondrial processes early after injury, it may be possible to alter the trajectory of neurodegeneration in vulnerable individuals.

## CRediT authorship contribution statement

Elika Z. Moallem contributed to writing the original draft, data analysis, and investigation. Hemendra J. Vekaria was involved in investigation and writing – review and editing. Teresa Macheda contributed to writing – review and editing and project administration. Kelly N. Roberts participated in investigation and writing – review and editing. Margaret R. Hawkins contributed to investigation and writing – review and editing. Samir P. Patel contributed to data analysis, interpretation, and writing – review and editing. Patrick G. Sullivan contributed to data analysis, interpretation, funding acquisition, and writing – review and editing. Adam D. Bachstetter was responsible for conceptualization, funding acquisition, project administration, writing the original draft, and data interpretation.

## Acknowledgments

Some figures in this manuscript were created using BioRender.com.

## Data availability

Data used in this study are available through the Open Data Commons for Traumatic Brain Injury (odc-tbi.org; RRID:SCR_021736), Moallem et al., (2025), https://doi.org/10.34945/F52W2X

## Declaration of generative AI usage in scientific writing

During the preparation of this work, the author(s) used ChatGPT and Claude to assist in evaluating the original text for clarity, completeness, and style. These tools provided editorial suggestions and recommendations; however, the author(s) reviewed and edited all content as necessary. The author(s) take full responsibility for the final content of the published article.

**Abbreviations**
Aß: amyloid beta
ABC: avidin-biotin complex
AD: Alzheimer’s disease
ANOVA: analysis of variance
AP: anterior-posterior
APP: amyloid precursor protein
APP/PS1 KI: amyloid precursor protein/ presenilin-1 knock-in
ATP: adenosine triphosphate
BCA: bicinchoninic acid
BSA: bovine serum albumin
CCI: controlled cortical impact
CHI: closed head injury
CO_2_: carbon dioxide
DAB: 3,3′-diaminobenzidine
EGTA: ethylene glycol-bis(β-aminoethyl ether)-N,N,N′,N′-tetraacetic acid
ETC: electron transport chain
F: female
FCCP: carbonyl cyanide-p-trifluoromethoxyphenylhydrazone
FDR: false discovery rate
HEPES: 4-(2-hydroxyethyl)-1-piperazineethanesulfonic acid
KCl: potassium chloride
KOH: potassium hydroxide
KH_2_PO_4_: potassium dihydrogen phosphate
M: male
MAD: median absolute deviation
ML: medial-lateral
MgCl_2_: magnesium chloride
NCLX: Na/Li/Ca exchanger
OCR: oxygen consumption rate
PBS: phosphate-buffered saline
PCA: principal component analysis
PDH: pyruvate dehydrogenase
PFA: paraformaldehyde
PPP: pentose phosphate pathway
PSI: pound-force per square inch
RCR: reactive oxygen species
SABV: sex as a biological variable
SEM: standard error of the mean
SD: standard deviation
TBI: Traumatic brain injury
WT: wild-type

